# Knowledge discovery with Bayesian Rule Learning for actionable biomedicine

**DOI:** 10.1101/785279

**Authors:** Jeya Balaji Balasubramanian, Kevin E. Kip, Steven E. Reis, Vanathi Gopalakrishnan

## Abstract

Biomarker discovery is critical for both biomedical research and for clinical diagnostic, prognostic, and therapeutic decision-making. They help improve our understanding of the underlying physiological processes within an individual. Discovery of biomarkers from complex biomedical datasets is done using data mining algorithms. Hundreds of thousands of biomarkers have been discovered and reported in literature but only a few dozen have been found to be clinically useful. This discrepancy is because statistical significance is not clinical relevance. Statistical significance only accounts for the correctness of the learned associations. Clinical relevance, in addition to statistical significance, also accounts for clinical utility such as cost-effectiveness, non-invasiveness, efficacy, and safety of the proposed biomarkers. We need models that are statistically significant and clinically relevant, all the while keeping it interpretable. Interpretable classifiers are more actionable in medicine because they offer human-readable explanations for their predictions. Traditional data mining methods cannot account for clinical relevance. We formulate this as a knowledge discovery problem. In computer science, knowledge discovery in databases is “a non-trivial process of the extraction of valid, novel, potentially useful, and ultimately understandable patterns in data”. Bayesian Rule Learning (BRL) finds an optimal Bayesian network to explain the training data and translates that into an interpretable rule model. In this paper, we extend BRL for knowledge discovery (BRL-KD) to enable BRL to incorporate a clinical utility function to learn models that are clinically more relevant. We demonstrate this using a real-world dataset to predict cardiovascular disease outcome. We evaluate predictive performance with the area under the receiver operating characteristic curve (AUROC) and clinical utility with the cost of the model. We show that BRL-KD successfully generates a set of models offering different trade-offs between AUROC and cost. Based on the clinical standard, a model with an acceptable trade-off can then be chosen.

## Introduction

Scientific investigators discover novel biomarkers for a wide range of purposes including improved clinical decision-making and guiding research directions in biomedicine. Novel biomarkers help improve our understanding of the physiological and pathophysiological processes being studied. Precise predictive biomarkers can help make accurate diagnostic, prognostic, and therapeutic decisions in clinical practice. A *biomarker* is defined as “a characteristic that is objectively measured and evaluated as an indicator of normal biological processes, pathogenic processes, or pharmacologic responses to a therapeutic intervention” [1]. Biomarkers commonly used in medical practice include macroscopic factors such as— age, smoking, blood pressure, gender, family history of a disease, environmental exposure, etc. They can also come from omics data, which includes molecular measurements from the DNA, RNA, proteins, metabolites, and epigenomic factors. Molecular biomarkers offer promising new tools for clinical decision making by providing insights into biological processes at an unprecedented resolution. Examples of molecular biomarkers include— blood cholesterol, blood glucose, harmful mutations in *BRCA* genes (a diagnostic biomarker associated with increased risk of developing breast or ovarian cancer) [2], increased serum prostate-specific antigen (a diagnostic biomarker associated with increased risk of prostatic disease) [3], etc.

*Biomarker discovery* is the process of discovering new biomarkers to improve our current understanding of the biological process under study. Biomarker discovery is typically data-driven and is performed using statistical data mining methods that find associations between candidate biomarkers (e.g., clinical factors, genes, transcripts, proteins, metabolites, etc.) and an outcome variable of interest (e.g., disease state, therapeutic response, etc.).

In 2016, Burke [4] reported that there were 768,000 papers indexed in PubMed directly related to biomarkers. While many of those papers claimed the markers to be clinically useful, very few molecular biomarkers are currently in clinical use today. Selleck at al. [5] further quantify this discrepancy with respect to cancer-related biomarkers— the European Society of Medical Oncology (ESMO) clinical practice guidelines for lung, breast, prostate, and colon cancers make weak recommendations for only about 20 molecular biomarkers. Clinical practice guidelines are considered a necessary minimum requirement for healthcare delivery. Therefore, it is apparent from the ESMO recommendations that molecular biomarkers have not yet achieved widespread clinical use.

The translation of discoveries made in basic biological research into medical practice is a slow process and is riddled with many challenges [6]. Selleck at al. [5] suggest that it is not enough to just identify variants but instead to identify *actionable* variants as biomarkers in order to revolutionize healthcare delivery. In order to improve translation of discovered biomarkers into medical practice, they call for a more focused biomarker discovery process using an evaluation criteria similar to the one set forth by Evaluation of Genomic Applications in Practice and Prevention Initiative [7] for evaluating genetic tests created for clinical and public health use. Their suggested four evaluation criteria are— 1) *analytical validity*, to test if the biomarker measurements are reproducible; 2) *clinical validity*, to validate if the biomarkers are indeed associated with the clinical outcome of interest; 3) *clinical utility*, to check if the biomarker will lead to improvements in clinical practice and eventually to public health; and 4) *other factors* such as cost-effectiveness, treatment or risk reduction, etc.

Shortliffe et al., [8] succinctly state this from an informatics standpoint— statistical significance is not clinical relevance. *Statistical significance* is the certainty that the discovered pattern— i.e., the discovered association between the biomarker and the clinical outcome— is correct. *Clinical relevance* is a measure of how useful the discovered pattern is in guiding clinical care. Clinical relevance, in addition to statistical significance, also accounts for clinical utility such as invasiveness, efficacy, safety, and cost associated with the discovered biomarkers. So, for example, if the newly discovered biomarker is just as precise but less invasive or cheaper than the current clinical standard, it has the same statistical significance but offers more clinical utility than the current clinical standard in practice.

Traditional data mining methods only focus on discovering statistically significant biomarkers. In this paper, we formulate the problem of identifying both statistically significant and clinically relevant models as a statistical knowledge discovery problem. Knowledge Discovery in Databases (KDD) is an important process of discovering useful knowledge from data. KDD is the non-trivial process of “identifying valid, novel, potentially useful, and ultimately understandable patterns in the data” [9]. In this definition, *patterns* refer to associations between biomarkers and clinical outcome of interest. They must be *understandable* to a clinician for it to be actionable in medical practice. *Valid* patterns generalize to unseen patient population. We can achieve this with models that contain patterns that are statistically significant. *Novel* biomarkers are useful in providing new research directions, whereas *useful* patterns can mean non-invasive, cost-effective, high efficacy, and safe biomarkers. In this paper, we will combine the terms *novel* and *useful* and just generally refer to both types as patterns with clinical utility.

High-throughput technologies are methods of automation for performing a large number of experiments in parallel. Molecular datasets for biomarker discovery are often measured using these technologies. Analyzing high-throughput datasets is challenging to data mining algorithms mainly because they are high-dimensional. High-dimensional datasets have significantly large number of candidate variables (e.g., clinical tests, genes, proteins, metabolites, etc.) but have relatively few examples (e.g., patient cohort) to help support any hypothesized association pattern between a biomarker and the clinical outcome. Any incorrectly found association is called a false positive. High-dimensional datasets considerably increase the risk of discovering false positives.

Bayesian Rule Learning (BRL) is a rule-based classifier generator that has been shown to be proficient in biomarker discovery from high-dimensional datasets and has been shown to perform better than state-of-the-art classifiers that are typically used for such applications [10,11]. This implies that BRL models generalize better to unseen data, which means they discover more *valid* patterns. Also, rule patterns are human-readable making them *understandable* patterns. For example, model reasoning using explicit propositional logical rules take the form *IF <condition>-THEN <consequent>*, which are easy for a human to read and understand. BRL also has additional utility in being able to incorporate prior domain knowledge [12], which can lead to an improvement in model predictive performance by helping further reduce the chance of discovering false positive associations and consequently generating more *valid* patterns. In many biomarker discovery tasks, we often have access to additional domain knowledge either from literature or curated bioinformatics databases that can help better tackle the challenges from the high-dimensionality of these datasets. Therefore, BRL currently learns *valid* and *understandable* patterns. To extend BRL to perform biomarker discovery, we must enable it to also discover patterns with higher clinical utility.

In this paper, we extend Bayesian Rule Learning (BRL) to perform knowledge discovery and call it Bayesian Rule Learning for Knowledge Discovery (BRL-KD). We hypothesize that BRL-KD can learn models containing biomarkers that, on average, have better statistical significance and better clinical relevance than BRL for the task of biomarker discovery. In the Materials and methods section, we describe the BRL algorithm, our implementation of BRL-KD algorithm, and finally the experimental design evaluating BRL-KD on a real-world diagnostic dataset on cardiovascular diseases to learn cost-effective diagnostic models. In the Results and Discussion section, we show the results from our experiment. In the Future Work section, we discuss the results and a few suggestions on other ways to learn clinical utility using BRL-KD, other than the task of learning cost-effective models. We then summarize our findings in the Conclusion section.

## Materials and methods

We first describe the BRL algorithm by introducing some notations used throughout the paper, describe the BRL model representation, the heuristic score it optimizes, and the search algorithm used to optimize the score. Next, we describe our extension of BRL to BRL-KD, specifically how we modified the heuristic score to include clinical relevance into the model learning process. Finally, in the experiment design, we use a real-world diagnostic dataset on cardiovascular diseases to demonstrate the BRL-KD model by evaluating it for— 1) statistical significance, with predictive performance using area under the receiver operating characteristic curve, and 2) clinical relevance, with cost-efficiency as an example, to help reduce the cost of the biomarkers selected by the classification model.

### Bayesian Rule Learning (BRL)

The Bayesian Rule Learning algorithm (BRL) takes a training dataset as input and searches over a space of constrained Bayesian belief-networks (BN) to identify the BN that optimally explains the training dataset. BRL uses a heuristic score, called the Bayesian score, to evaluate a candidate BN proposed by the search algorithm. After finding the optimal BN model that maximizes the heuristic score, BRL infers a rule model from the optimal BN. The constrained Bayesian network structure and its representation is described in the next subsection. We then describe the heuristic Bayesian score, and finally the greedy best-first search used to find the BN model that optimizes the Bayesian score.

#### Model representation

We denote the input training dataset as *D*. Dataset *D* contains a set of *n* discrete-valued, candidate predictive variables *X*_1:*n*_. These are the *n* candidate biomarkers measured during the study that generated the data. We use a subscript to represent the *i*-th variable, *X_i_*. Additionally, *D* also contains a specified discrete target variable, *Y*. This is the clinical outcome of interest for this data analysis task, for example, disease state, *Y* = {*Case, Normal*}. So, the dataset *D* = {*X*_1:*n*_, *Y*}. The database as m observations or instances (e.g., number of individuals in the study cohort). We refer to the t-th instance as 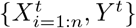.

A generalized Bayesian network is a probabilistic graphical model that uses a directed acyclic graph (DAG) to represent the full joint probability distribution of the variables in a problem domain [13]. A Bayesian network, *B*, is a tuple consisted of a DAG network structure *B_S_* and a set of parameters Θ i.e., *B* = (*B_S_*, Θ). The network structure consists of nodes and edges. A node is a variable in the domain and edges represent probabilistic dependencies between the variables. Variable *X_j_* is considered a parent of *X_i_* if there is a directed edge from *X_j_* → *X_i_*. This implies that variable *X_i_* is probabilistically dependent upon *X_j_*. A set of parents of variable *X_i_* is represented as Π*X_i_*. Each node, *X_i_*, in the network contains a parameter, *θ_i_*, which is a probability distribution of the values taken by the node conditioned on its parents, *θ_i_* = *p*(*X_i_*|Π*_X_i__*). This information from the parent is sufficient because each variable is independent of its non-descendants, given its parents [13]. Parameter set Θ is just the collection of the probability distributions of all the nodes of the network. Bayesian networks provide considerable computational savings by representing a full joint probability distribution by decomposing the probability distribution of each variable to be dependent only on its parents. For a brief tutorial on generalized Bayesian networks, please refer to Heckerman [14].

In this paper, we will only look at a special case of a generalized Bayesian network, called *constrained Bayesian network* (BN). A BN only consists of the target variable, *Y*, and its parents Π_*Y*_. We illustrate this using Fig 1.

**Fig 1.**
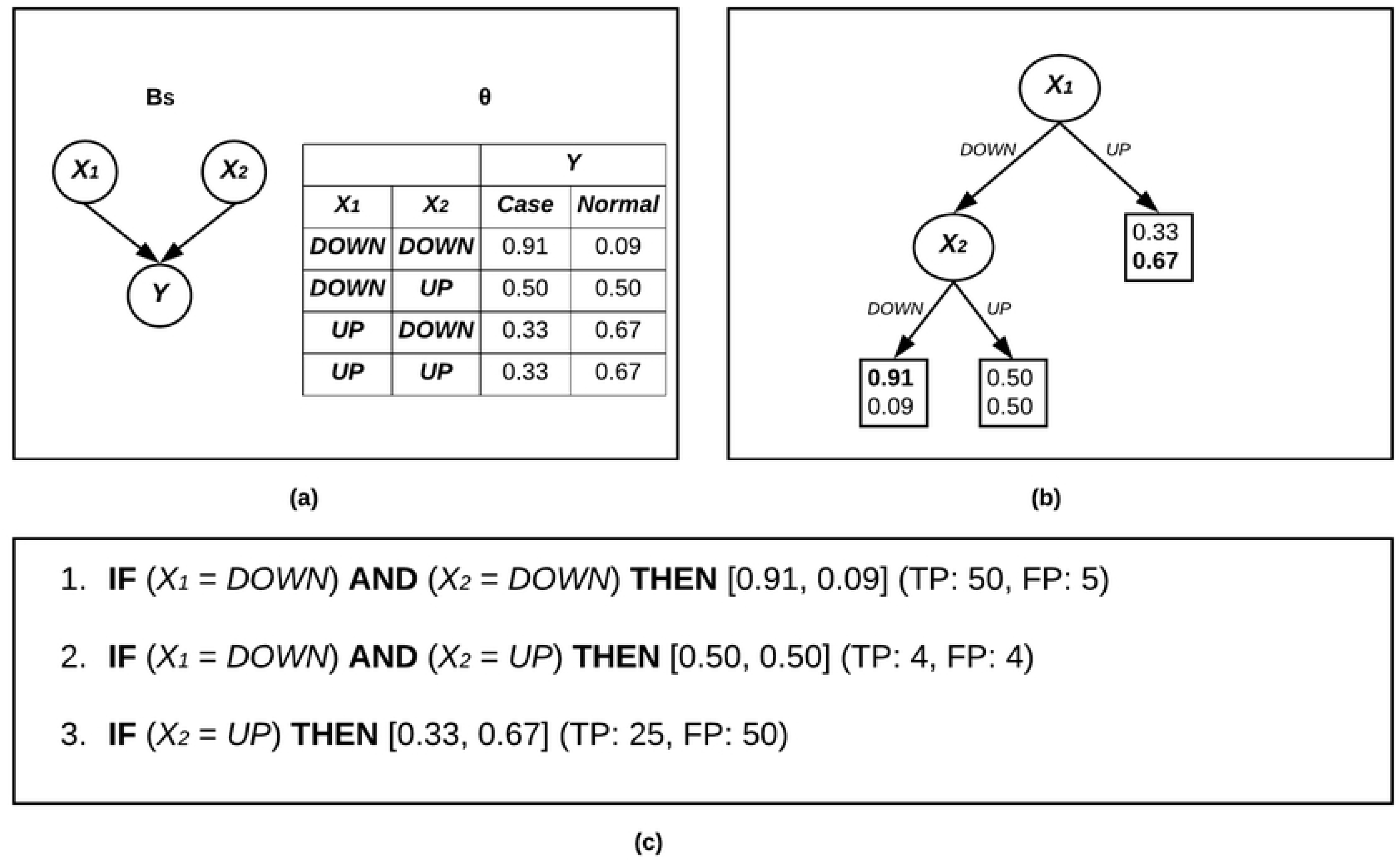
Bayesian Rule Learning (BRL). BRL representation. (a) The constrained Bayesian network structure and parameter associated with the target variable. (b) The parameter represented as a decision tree. (c) BRL translating the decision tree into rules.

Fig 1a, shows the BN *B_S_* and the parameter associated with the target variable *θ* = *p*(*Y*|*π_Y_*). The BN here contains the target variable *Y* and it has two parents Π_*Y*_ = {*X*_1_, *X*_2_}. The parameter *θ* contains all possible variable-value pairs of the parents of *Y*. Each variable-value assignment of its parents is called a *configuration*. For each configuration, the probability distribution over values of *Y* is shown in the table. This table is called the conditional probability table or CPT. The CPT is represented as a decision tree in Fig 1b. To understand the properties of this representation, please refer to our earlier work [11]. Eventually, BRL translated this decision tree into IF-THEN rules as shown in Fig 1c. Each rule is a path from the root to leaf of the decision tree representation. For example, the first rule is read as follows— if the variable (biomarker) *A* takes the value *DOWN* (is under expressed) and the variable *B* also takes the value *DOWN*, then it implies that the target variable (disease state) takes the value *Case* with a probability 0.91 and takes the value *Normal* with a probability 0.09. This rule is supported by 50 true positive (*TP*) examples i.e., the number of instances in D, where both the left-hand-side and the right-hand-side of the rule is true. The rule also matches 5 false positive (*FP*) examples i.e., the number of instances in D, where only the left-hand-side of the rule is true but the right-hand-side of the rule is false. How we compute the probability associated with each target value is shown in the end of the next subsection.

#### Heuristic score

To evaluate the quality of the BNs found during the search, we need a heuristic score to evaluate how well it explains the input training data. Buntine [15], under certain assumptions, proposed a heuristic Bayesian score called the BDeu (Bayesian Dirichlet equivalence uniform) score to compute the joint probability of a generalized Bayesian network and the input training data. We modify this score to only evaluate for the constrained Bayesian network containing the target variable and its parents. This modified score is shown in Eq 1.

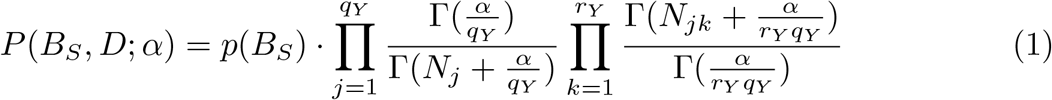

Here, index *j* iterates though each of the *q_Y_* possible variable-value configurations of the parents of the target *Y*. Index *k* iterates through each value taken by the target variable *Y*. The hyperparameter *α*, is called the *prior equivalent sample size*. It expresses the strength of our belief in the prior distribution over the networks. The expression 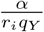 assigns a uniform prior probability to each parental configuration. And, 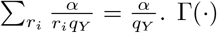 is the gamma function, i.e., Γ(*x* + 1) = *x*!. *N_ijk_* is the number of instances in *D*, where the target variable *Y* takes the value *k*, while its parents are in configuration specified by *j*. And, *N_ij_* = Σ_*k*_ *N_ijk_*. All the terms described so far is collectively called the likelihood function.

The first term *p*(*B_S_*), in Eq 1 is the prior term. It is the prior distribution that represents our prior belief about which of the models in the model space is likely to be correct, before we saw the training data. In our earlier work [12], we showed how to use this term to incorporate prior domain knowledge from literature to help improve the model predictive performance. In this paper, however, we will use it to constrain the search to prefer models with fewer parents added to the target variable. This makes an assumption that models with fewer variables are more likely to be correct. This can be helpful while modeling high-dimensional data as the search would try to reduce false positive discoveries.

To achieve sparser models, we use a prior distribution score as described by Koller et al. [16] to prefer models with fewer parents added to any given node in the network. This is prior term is shown in Eq 2.

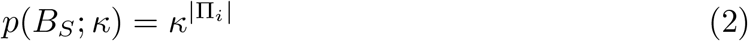

Here, the exponent term |Π_*i*_| is the total number of parents that the target variable has in structure *B_S_*. Hyperparameter *κ* is a value ranging between 0 < *κ* ≤ 1. A value of *κ* =1 represents a uniform prior over all network structures. While, smaller values of *κ* generates a distribution that prefers structures with nodes having fewer and fewer parents, larger values for *κ* generates a distribution that allows more and more parents.

We combine Eq 1 and Eq 2 to obtain the heuristic score for BRL as shown in Eq 3.

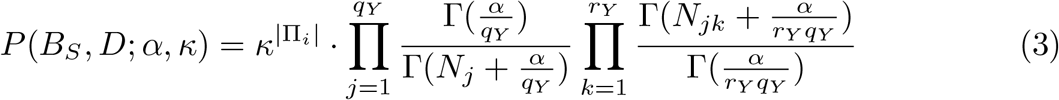

Once we have the optimal BN, we can predict the target variable value of an unseen test instance, *Y^t^*, given the value of its parent variables, *X^t^*, the learned optimal BN, *B_S_*, and the training data *D*. We compute this using the expectation of the parameter associated with the target variable as shown in Eq 4.

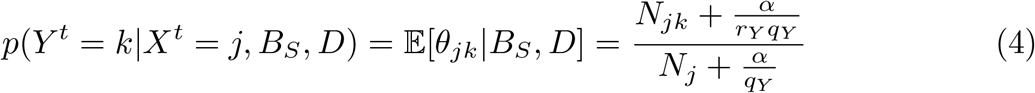

To make a prediction, we simply return that *k* value for which the above value is the highest i.e., arg max_*k*_*p*(*Y^t^* = *k*|*X^t^* = *j, B_S_, D*).

We now need a search algorithm to find the BN that maximizes this score. We describe this search algorithm in the next subsection.

#### Greedy best-first search

In this paper, BRL uses a greedy best-first search to find the BN that maximizes the heuristic score in Eq 3. The greedy best-first search algorithm described here is based on the same as the one described in our earlier work (see [11]). We modify it slightly to remove the user parameter, *MAX_CONJ*, from the search. The constraint in the complexity of the model, in this paper, comes from the prior term we specified in the heuristic score in Eq 3.

The BRL search algorithm takes as input, the training dataset *D*. In this paper, we set the *α* = 1, similar to how we did in the previous paper [11]. We set the value of *κ* = 0.01. This choice is a small value, chosen after having run a few experiments with simulated datasets. In this paper, we do not concern ourselves with optimizing the prior term, so we keep it fixed.

In the first step of the search algorithm, BRL initializes with *n* BNs, each containing the target variable Y and a unique parent variable selected from *X*_*i*=1:*n*_. Each BN is evaluated by the heuristic score in Eq 3. The BN with the highest score is selected for the next iteration of the algorithm. Let’s call this model, *prev.best*, i.e., the model that did the best in the previous iteration. In each subsequent iteration, a new parent is added to Y, such that it is not already present in its parent set Π_*Y*_. After having tried each variable to its parent set, one at a time, the BNs are evaluated again using the heuristic score. If the score of any of these models improve upon the score of the model which performed the best in the previous iteration, *prev_est*, then the new specialized model is chosen as the best model in this iteration. This iteration continues, until adding another variable from *D* does not help improve the heuristic score of the BN. This model is returned by BRL as the optimal BN.

BRL then translates the BN into a set of rules as shown in Fig 1. For a more detailed explanation of this search algorithm, please refer to Lustgarten et al. [11].

### Bayesian Rule Learning for Knowledge Discovery (BRL-KD)

As mentioned in the introduction section, BRL helps us find *valid* and *understandable* rule patterns from the training data. However, it does not account for clinical utility. Let us re-visit the heuristic score in BRL, as shown in Eq 3. Here, the structure prior term is *p*(*B_S_*) = κ^|Π_*i*_|^, is the prior distribution that represents our belief in which of the models in the hypothesis space is likely to be correct *a priori*. We set the value to prefer models with fewer parents. In our previous work, we developed a method called BRL_p_ [12], we used the structure prior term, *p*(*B_S_*), to incorporate prior domain knowledge to create a bias in the search to prefer network substructures that have been shown to be promising in the literature. The rest of the term in the equation is called the likelihood function that encodes the likelihood that the observed training data was generated by a given BN model. The likelihood function gives us a measure of how well the model fits the data. Generally speaking, the better the model fits the data, the more likely it is to generalize to unseen test data. The prior term encodes the prior probability of which of the BNs is the correct data-generating model. Together, the likelihood function and the prior term assist the search algorithm in identifying promising candidate BNs that are most likely to have generated the training dataset. In KDD definition, these two terms help the search algorithm find *valid* patterns from data.

To bias the search to prefer more *useful* patterns, we modify the heuristic score to Eq 5.

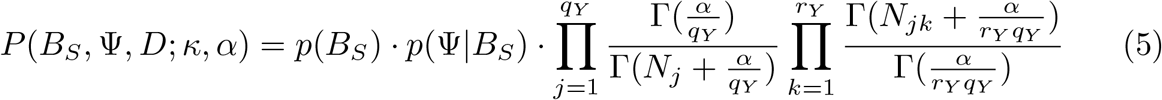

This equation encodes the joint probability of the BN structure (*B_S_*), the data (*D*), and the clinical relevance as encoded by the utility function (Ψ). The utility function *p*(Ψ|*B_S_*) takes the BN structure as input and outputs the clinical relevance of the model. The term *p*(Ψ|*B_S_*) encodes a probability distribution that represents our belief about which of the models in the hypothesis space is more clinically relevant.

We encode the utility function similar to how we encoded the prior distribution over models in BRL_p_. We use the informative prior as specified by Mukherjee et al. [17]. The utility function is a log-linear combination of weighted real-valued function (concordance function) of the network structure, *B_S_*. The utility function monotonically increases with the increase in clinical relevance of the model. The utility function is shown in Eq 6.

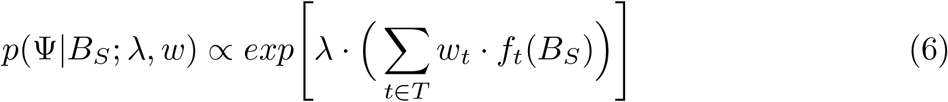

Let us see an example to understand how to use this utility function. Say, the current clinical standard being used in medical practice is expensive and we want cheaper yet still effective models to replace them. In other words, we want more cost-effective models. The utility function is encoded to score cheaper models, higher. One way to compute the model cost is by adding the cost associated with each biomarker in the model. To enable BRL-KD to look for cost-effective models, the set *T* in Equation 6 iterates through each biomarker in the dataset. Weight wt is the weight of the biomarker *t*. For cost-effectiveness, we set the weight to the cost of the biomarker. The concordance function here can be an indicator function that returns 1, when the variable *t* exists in the BN structure, *B_S_*. We use the same scoring function, we have described here, to demonstrate BRL-KD on a real-world dataset in the next subsection.

An important consideration while using this utility function is to encode it as a maximization problem. This is because we maximize the heuristic score in the greedy best-first BRL search. However, currently we have encoded the current utility function in such a way that we want to minimize the function (i.e., minimize the total cost). To keep this consistent with the Bayesian score heuristic, we must convert the utility function into a maximization problem. We can do that by simply encoding the negative value of cost, instead of the cost. Now, the utility function needs to be maximized in order to minimize the overall cost.

Another important consideration while using the utility function is the value set for the weights. When encoding the cost, some markers may cost a few US dollars, some others may cost thousands of dollars. This will lead to this term being either too large or too small as a result of it being on the exponent term of the utility function. To avoid this, we must scale this value. For example, we can perform min-max scaling for the values to range between 0 and 1.

Let us now try to understand the role of the λ hyperparameter. In Eq 6, λ = 0 signifies no confidence in specified values of each biomarker. When we set this value, BRL-KD behaves exactly like BRL. A large value of λ ≥ 25 asks the search algorithm to prioritize cost before looking at the likelihood function. Here, BRL-KD picks the cheapest variable available at each iteration and if more than one variable are similarly priced, BRL-KD then consults the likelihood function to break the tie. The user may optimize over a range of values for λ in between 0 and 25. The set of models generated by varying λ generates a set of solutions (called a Pareto set), accommodating our cost constraints while incorporating our specified knowledge at varying degrees. We can leave the decision of which model to use based on the circumstances at the point-of-care. One way to cut-down on the number of models to examine further is by specifying a constraint like the required AUROC (or more complex metrics like cost per life years gained) achieved by each λ over 10-fold cross-validation.

BRL-KD maximizes the heuristic function in Eq 5 to identify patterns that are *valid, novel, potentially useful*, and *understandable*, from the data. To specifically seek novel patterns, we may alter the utility function to encode a metric that quantifies how well known it is in literature to have each of the biomarkers to be associated with the disease. We explain this situation further in the Future Work section. For now, let us design and observe BRK-KD in practice using a real-world example.

### Experiment design

The goal of the experiment here is to study the changes in behavior of BRL-KD models as a result of altering the hyperparameter λ. Specifically, we want to observe the changes in two evaluation metrics— 1) a clinical relevance metric, here we calculate the total cost of all the biomarkers selected by the model, and 2) a predictive performance metric, here we use area under the receiver operating characteristic curve (AUROC).

In this experiment, we use a real-world dataset collected to learn differential features between individuals who are at risk to develop cardiovascular disease (CVD) and those who are unlikely to develop the disease in the near future. Here, we assume that the most cost-effective diagnostic model is the clinically most relevant model. Cost-effective models have both good predictive performance and are cheaper than clinical standard currently used in practice.

#### Data collection and pre-processing

Heart SCORE (**S**trategies **C**oncentrating **O**n **R**isk **E**valuation) is an ongoing longitudinal prospective study, initiated in 2003, that follows 2000 middle-aged, primarily Caucasian and African-American individuals from Allegheny County, Pennsylvania, USA [18]. We consider two types of variables measured from individuals in the study at the time of enrollment. They are— clinical and metabolic variables.

The clinical variables include— age, sex, race, patient medical history, physical examination, medications, Vertical Auto Profile test for lipids, finger stick tests, and questionnaires about the individual’s physical activities, lifestyle markers, social network, diet, sleep quality, and various psychological questionnaires. The data has a total of 654 such clinical variables. The metabolic variables had 1228 biochemicals including 893 named and 335 unnamed biochemicals. These biochemicals include xenobiotics, co-factors, vitamins, and metabolites from amino acid, lipid, carbohydrate, and energy metabolism. Out of the 2000 study participants, only 1901 had metabolites measured for them. So, the dataset had 1901 instances.

The data had to be pre-processed to prepare it for analysis using BRL. We first had to clean the data. The dataset had 608 clinical variables. For this data analysis, we assume that the missing values are Missing Completely at Random (MCAR). Under the MCAR assumption, when the variable with missing values have ≤ 5% of the values missing, we can impute a single complete dataset, with minimal bias, using median/mode value imputation. In this analysis, we dropped the variables with > 5% of its values missing. The rest of the variables had missing values imputed with median/mode imputation.

We define the outcome variable of interest as Major Adverse Cardiac Event or MACE, which includes any of— Cardiac death, myocardial infarction (MI), acute ischemic syndrome (AIS), or revascularization.

The final dataset had 1617 candidate predictive variables including 389 clinical variables and 1228 metabolic variables. There are 1901 instances with 101 positive cases of individuals who eventually developed a MACE outcome by 2018. The remaining 1800 individuals were labeled negative.

We randomly split the dataset into 70% training data and 30% test data. We will use the training data to identify our preferred value for hyperparameter λ by observing the average cost and average AUROC over cross-validation, achieved by a set value for λ. The average of the metrics is computed over 10-fold cross-validation. We then choose the λ with our preferred trade-off between cost and AUROC, and use that to model the training dataset. We observe these models and evaluate their performance on the 30% held-out test set.

Since BRL-KD can only handle discrete valued data, we discretize the continuous valued variables from the Heart SCORE data under cross-validation. When the training data is split into 10-folds, in each iteration of the cross-validation, we discretize on the training-fold using the supervised minimum description length (MDL) principle method [19] and use the learned bins to discretize the test-fold dataset. Once we determine the ideal value for λ from the cross-validation experiment, we discretize the 70% training data using the MDL method and use the learned bins to discretize the 30% test data.

#### Methods compared

We will study the change in behavior of BRL-KD with the change in hyperparameter λ. We encode the term *p*(Ψ|*B_S_*) in Eq 6 with a distribution that represents our belief about which of the models in the hypothesis space is more cost-efficient. We do this using Eq 7.

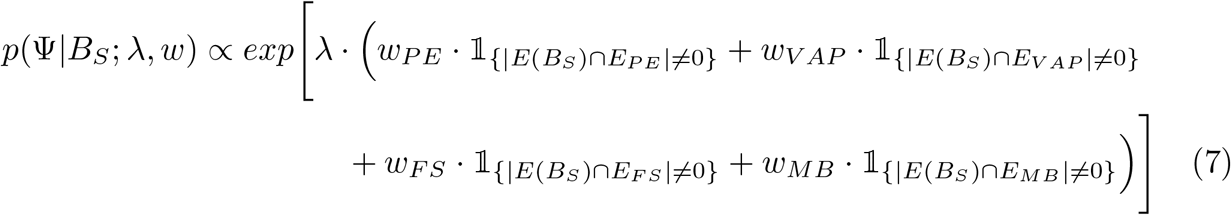

In this equation, λ represents the relative importance of considering cost as opposed to the likelihood term (that tries to optimize predictive performance). The set of weights, *w* = {*w_PE_, w_VAP_, w_FS_*, *w_MB_* }, represent the cost of physical examination (*w_PE_*), cost of Vertical Auto Profile test for lipids (*w_VAP_*), cost of finger stick tests (*w_FS_*), and the cost of running a full metabolome profile of 1228 biochemicals (*w_MB_*).

We want to state a disclaimer that none of the values encoded in this experiment represents reality and were not done consulting a medical practitioner. They were merely estimated to demonstrate the functionality and behavior of BRL-KD as a proof of concept. The numbers in the next paragraph were obtained using simple online search queries. In reality, the cost can be influenced by many factors including time, insurance, medicare, and the location of point-of-care. For now, we assume that the cost associated with each biomarker presented in the next paragraph is correct and we observe the function of BRL-KD in light of having specified such values in practice.

The Heart SCORE dataset contains clinical and metabolic variables. For all the demographic and questionnaire-based variables, we set the cost of the biomarkers to $0. We assume that it is free to obtain markers that can be supplied by asking the individual or can be obtained from their medical record. We set the cost of a physical exam, *w_PE_* = $146. By setting the value of *w_PE_* = 146, we state that if the BRL model requires the use of any one or more variables from the physical examination report (e.g., height, weight, or body mass index), the model would incur a cost of $146. The cost of Vertical Auto Profile (VAP) test for lipids is set to *w_VAP_* = $2813, for finger stick tests *w_FS_* = $1, and the whole metabolic panel *w_MB_* = $1000. We re-emphasize that the cost in this example is purely hypothetical and instead the focus of experiments in this paper is to demonstrate and evaluate the mechanism of being able to incorporate such information into the model learning process. For example, our cost estimate for VAP test might seem unrealistically steep. This estimate was based on very detailed quantification of lipid subfractions and profiles, and is on par with other expensive tests, such as imaging.

The costs range in thousands leading to this probability ranging widely. To prevent this, we scale the cost using min-max scaling to have the cost values between the range 0 and 1. We also take the negative value of the scaled cost in order to convert this utility function into a maximization problem.

#### Evaluation metrics

We measure two metrics— 1) clinical relevance using the total cost of the model, and 2) predictive performance using AUROC.

The cost metric is simply the sum of the costs of each marker selected by the BRL classifier. If the BRL model would select any variable from one of the sets with specified costs, the model would incur a cost as specified by the associated weight. For example, if the BRL model selects 2 metabolic variables, it would still only incur the cost of *w_MB_* once, since we assume that the whole metabolic panel is run. Again, we emphasize that our description of utility is meant to merely depict a real-world application and that our choice may not reflect reality. Our goal is to evaluate the use of BRL-KD in a possible real-world scenario.

## Results and Discussions

### Performance over cross-validation

On the 70% training data, we perform 10-fold cross validation. In each fold, the training data is split into 90% train and 10% development data. The 90% train is used to learn a BRL-KD model. We try different values of the hyperparameter λ = {0,1, 2, 4, 6, 8,10,12,14,16,18, 20}. We plot the average cost and average AUROC across 10-fold cross-validation in Fig 2.

**Fig 2.**
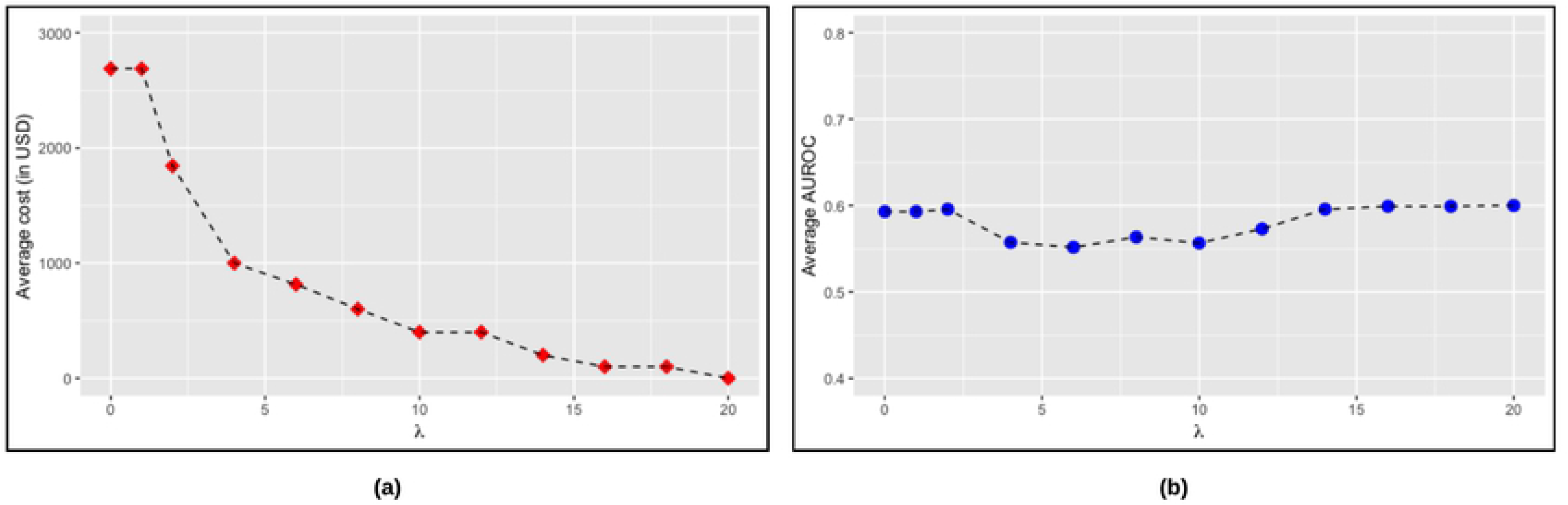
BRL-KD evaluation over cross-validation. Average cost and AUROC, over 10 folds, of the models learned under different values of hyperparameter λ. (a) Change in average cost. (b) Change in average AUROC.

We observe that the average cost steadily decreases with the increase in the value of λ. The average AUROC is more or less steady with the change in λ. The value of λ = 0 sets the whole expression of *p*(Ψ|*B_S_*) in Eq 5 to 1. As a result, this heuristic corresponds to that of plain BRL. The average cost of this model is $2687.8 and the associated average AUROC is 0.5930.

As we increase the value of λ, the BRL-KD model starts to pick more cost-efficient markers. This means markers that are of similar or slightly poorer quality but at a cheaper cost. As we increase the λ value, we get cheaper and cheaper BRL-KD models. Value λ = 4 corresponds to an average cost of $1000 and average AUROC of 0.5575. This λ value presents with a cheaper option at the loss of some predictive power. Value λ =10 corresponds to an average cost of $400.2 and average AUROC of 0.5566. Value λ = 20 corresponds to an average cost of just $0.2 and average AUROC of 0.6001. This λ value results suggest that there are similarly good AUROC performing models by simply using variables that are available for free (according to our definition).

### Performance over held-out test set

We now look at the models learned on the overall 70% training dataset, for λ = {0,4,10}, to observe the BRL-KD models. Note that we also looked at models with greater values of λ but those models remained consistent after λ = 10. The value λ > 10 corresponded with models with a total cost of $0 so there was nothing to optimize further using the λ hyperparameter. We estimate the AUROC by evaluating the model on the held-out 30% test dataset. Together, the set of BRL-KD models, offering trade-offs between cost and AUROC, is called a Pareto set of solutions.

The resulting BRL-KD model cost and AUROC is plot in Fig 3. We see that with increasing λ, we get cheaper models but we get them at a loss of AUROC performance on the test set.

We now observe the BRL-KD models generated by the three values of λ. Fig 4a, shows the BRL-KD rule model we get when λ = 0. This sets the whole expression of *p*(Ψ|*B_S_*) in Equation 5 to 1. This means no attempt is made to search for cost-efficient markers in the dataset and is equivalent of just running plain BRL. This model picks up a metabolic variable *N2,N2-dimethylguanosine* (cost = $1000), a variable from the VAP test *HDL subfraction 2A* (cost = $2813), and a demographic variable *Age* (cost = $0). So, the total cost of this baseline model is $3813 and the AUROC performance of this model on the held-out test set is 0.6906.

**Fig 3.**
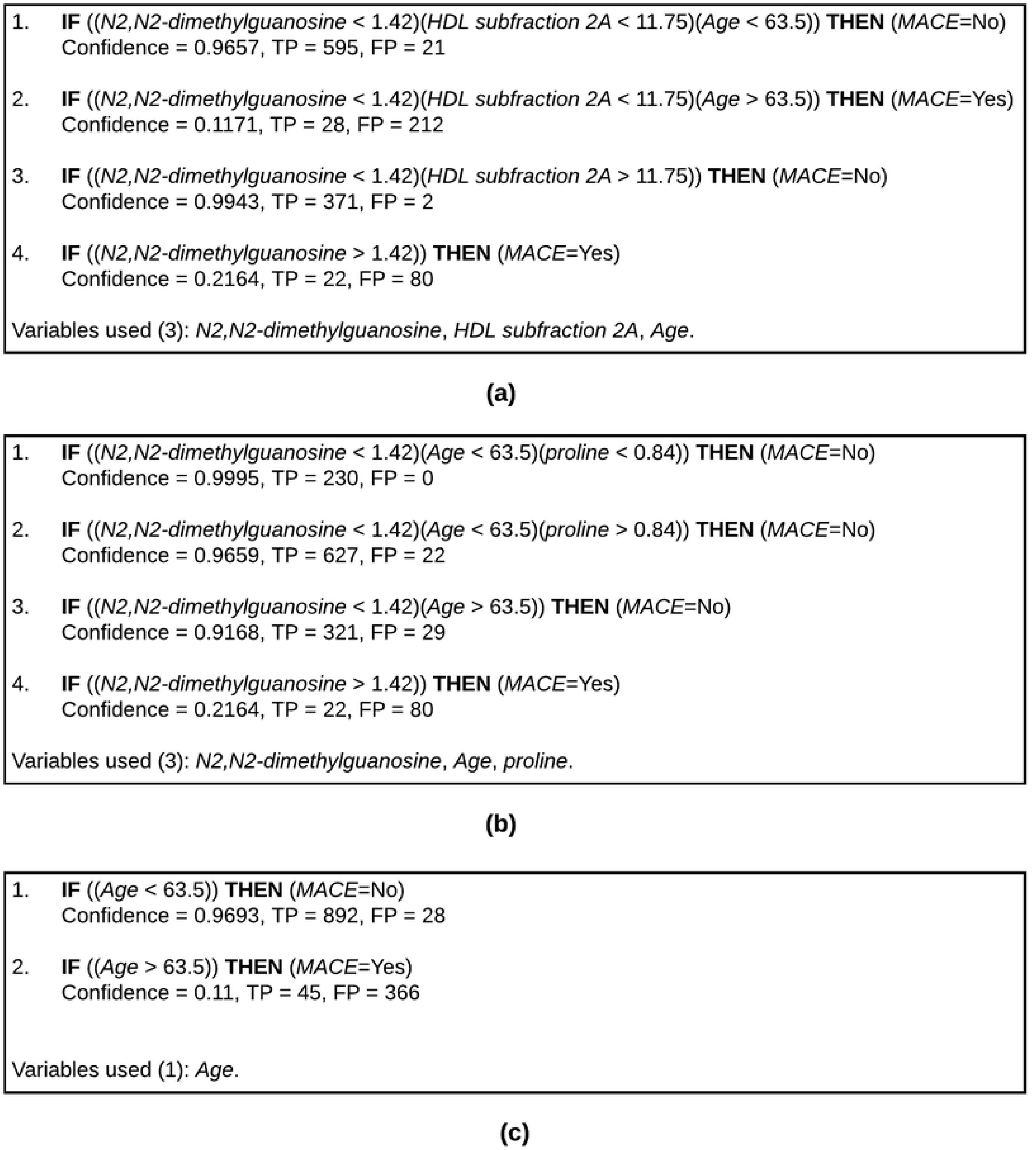
BRL-KD evaluation over held-out test set. Cost and AUROC on the test set for various λ values.

**Fig 4.**
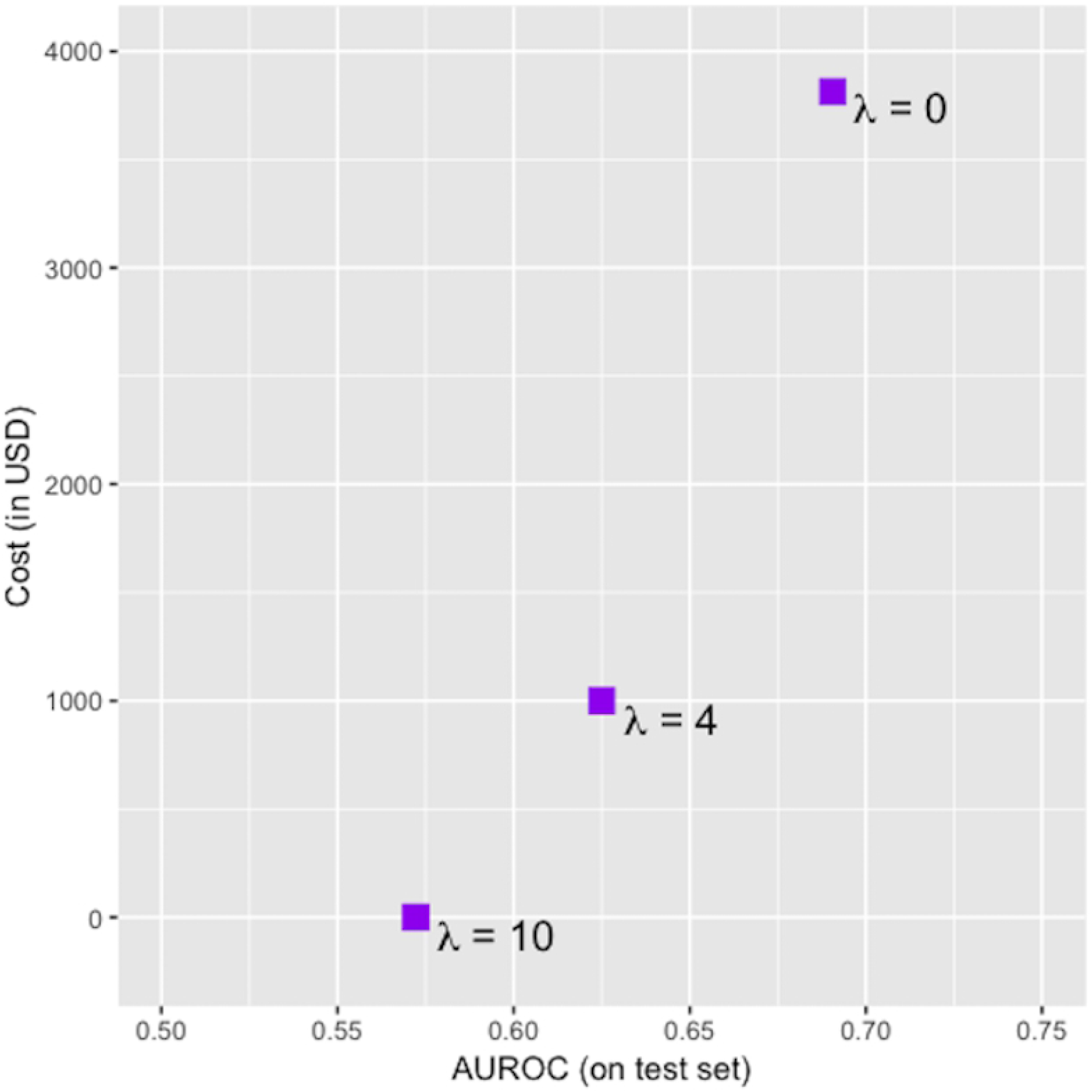
BRL-KD models. BRL-KD models learned from various λ values.

Fig 4b, shows the BRL-KD rule model we get when λ = 4. This model picks up two metabolic variables *N2,N2-dimethylguanosine* and *proline* (cost = $1000) and a demographic variable *Age* (cost = $0). So, the total cost of this cost efficient model is brought down to $1000 and the AUROC on the test set is 0.6250. Trying to come up with more cost efficient markers, BRL-KD loses the most expensive marker from the VAP test and substitutes it with a metabolic variable. However, we see that this also leads to a loss of AUROC on the test set.

Fig 4c, shows the BRL-KD rule model we get when λ = 10. This model uses a single demographic variable that is available for free, *Age*. This leads to a further saving of cost down to $0. However, we do that by compensating on the AUROC performance of 0.5722.

### BRL-KD compared to other rule-based classifiers

We additionally learned three popular rule-based classifiers from the Weka machine learning suite (version 3.9.3) [20], namely— C4.5 [21], a decision tree learning algorithm (where each path from root to leaf in the tree is translated into a rule), RIPPER [22], and PART [23].

C4.5 attains an AUROC of 0.4790 on test set and incurs a cost of $3813. But the C4.5 model requires 12 variables, including 8 metabolites and 17 rules (number of leaves in the decision tree). RIPPER obtains an AUROC of 0.4945 on the test set but only costs $1000. The AUROCs of C4.5 and RIPPER indicates a random classifier and are therefore, uninformative. RIPPER also requires 13 variables, including 9 metabolites and needs 4 rules. Comparatively, PART performs better with an AUROC of 0.7120 but costs $3959, more than any model seen so far. This is because PART requires 44 variables and a total of 31 rules.

BRL, on the other hand is parsimonious, requiring only 3 variables for an AUROC of 0. 6906 and costs $3813. To make a decision on weather or not we want to use any of the BRL-KD models, we must compare to the current clinical standard. The BRL-KD model enables us to know if there exists a model with a better or similar predictive performance at a cheaper cost.

## Future Work

The goal of this paper was only to demonstrate and evaluate the functionality of BRL-KD. We used a real-world dataset to do this and used an arbitrarily defined utility function. The models developed in this paper do not perform well enough to be considered for clinical use. However, this paper provides motivation to try and use the functionality of BRL-KD to explore real-world datasets with a reasonably accurately defined clinical utility function.

In this paper, we only considered one definition of clinical utility, i.e., cost-effectiveness. However, clinical utility can also mean novelty, non-invasiveness, specificity, etc. Or any combination of those definitions. An important future work would be to define new ways to encode clinical utility function using the same general form as shown in Eq 6. To do this, we must first try to quantify this utility metric at a biomarker level. For example, say we are interested in finding models containing novel biomarkers. One way to quantify novelty is by finding the number of references in biomedical literature that associate each candidate biomarker with the disease. This can be done using a test mining tool like KinderMiner [25]. Similar metric was quantified by Kleiman et al. to discover novel lab tests [24]. This metric can be used to encode the utility function in BRK-KD to return models with novel biomarkers. These models are clinically useful by providing new biomedical research directions.

Similarly, we can encode biomarker specificity by looking at gene-disease ontologies and encoding the weights with the reciprocal of the number of known diseases associated with each input gene. By maximizing such a function, BRL-KD would prefer models containing biomarker with fewer known associated disease. These models are clinically useful by providing tests that are specific to only the disease under study and not a biomarker that indicates many associated illnesses.

## Conclusion

The goal of the experiments was to demonstrate the functionality and behavior of BRL-KD and not to actually find a workable clinical model for CVD diagnosis. We saw that increasing the λ hyperparameter helped us specify a range of possible trade-off values between clinical relevance and predictive performance. We used cost as the utility function measuring clinical relevance. Using BRL-KD, we tuned the hyperparameter λ to study a range of different trade-offs between cost and predictive performance.

To pass the four evaluation criteria posited by Selleck at al. [5], as described in the Introduction section, we need to consider everything right from the experimental design used to collect the dataset to the statistical data analysis method. However, this paper only dealt with the statistical data analysis step. We assumed that the candidate biomarkers measurements, chosen for data collection in the study, were reliable and reproducible. However, to improve the chances of successful translation of discovered biomarkers into clinical practice, the scientific investigators must also consider efficient experimental design for data collection.

The ultimate decision on λ requires us to know a few things about the current clinical standard being used in practice. This will help us determine from the data if there exists a λ value that gives us a model that is clinically more relevant than the one being used in medical practice today. From the experiments in this paper, however, it appears to be clear that BRL-KD is a useful tool to help us find clinically more relevant models and encourages usage in a real-world problem.

The Java code for BRL-KD has been made open source and is available for public use from the GitHub repository (https://github.com/jeya-pitt/Bayesian-Rule-Learning).

## Acknowledgments

The authors gratefully acknowledge past grants from the National Library of Medicine, Award Number R01LM010950, National Institute of General Medical Sciences, Award Number R01GM100387 and an award from the Pittsburgh Health Data Alliance to VG that enabled some of the research infrastructure for this work. The content is solely the responsibility of the authors and does not necessarily represent the official views of the National Institutes of Health. All authors declare no conflicts of interest.

